# Exploring the adhesion properties of extracellular vesicles for functional assays

**DOI:** 10.1101/2024.10.28.620659

**Authors:** Bianca C. Pachane, Bess Carlson, Suzanne E. Queen, Heloisa S. Selistre-de-Araujo, Kenneth W. Witwer

## Abstract

The “stickiness” of extracellular vesicles (EVs) can pose challenges for EV processing and storage, but adhesive properties may also be exploited to immobilize EVs directly on surfaces for various measurement techniques, including super-resolution microscopy. Direct adhesion to surfaces may allow examination of broader populations of EVs than molecular affinity approaches, which can also involve specialized, expensive affinity reagents. Here, we report on the interaction of EVs with borosilicate glass and quartz coverslips and on the effects of pre-coating coverslips with poly-L-lysine (PLL), a reagent commonly used to facilitate interactions between negatively charged surfaces of cells and amorphous surfaces. Additionally, we compared two mounting media conditions for super-resolution microscopy (SRM) imaging and used immobilized EVs for a B-cell interaction test. Our findings suggest that borosilicate glass coverslips immobilize EVs better than quartz glass coverslips. We also found that PLL is not strictly required for EV retention but contributes to the uniform distribution of EVs on borosilicate glass coverslips. Overall, these findings suggest that standard lab materials like borosilicate glass coverslips, with or without PLL, can be effectively used for the immobilization of EVs in specific imaging techniques.

## Introduction

Extracellular vesicles (EV) adhesive properties arise from phenomena including charge (the EV surface is usually negatively charged) and molecular affinities due to the presence of, e.g., integrins and other adhesion molecules (1). Multiple studies have investigated EV interactions with specific materials, often in hopes of reducing negative consequences such as loss to surfaces during EV storage (2–4). However, direct immobilization of EVs onto materials such as glass might also allow for the improvement of techniques including fluorescence microscopy.

Traditionally, EVs have been captured for imaging using molecular affinity reagents such as antibodies (5,6). Alternatively, glass slides and coverslips can also be coated with poly-L-lysine (PLL) or poly-D-lysine (PDL). As polymerized amino acids with cationic charge, these reagents enhance the capture of negatively charged particles (7,8).

Non-specific immobilization may be especially valuable now that super-resolution (SRM) has evolved to enable semi-quantitative multiplexed single-molecule characterization of the composition of single EVs and thus convey a richer sense of EV heterogeneity (9). For this reason, we investigated several aspects of EV immobilization onto surfaces for imaging. We tested the direct adherence of EVs onto borosilicate glass and quartz coverslips, further probing the extent to which pre-coating with PLL enhances EV immobilization. We also examined mounting conditions for SRM imaging and determined that glass-immobilized EVs could also be used for cell interaction assays. We submit that these findings will be valuable for other investigators who wish to image immobilized EVs.

## Materials and Methods

### EV production: cell culture and transfection and EV separation

Using the Expi293™ expression system (Gibco #A14635), Expi293F cells were grown and transfected with the pLenti-palmGRET reporter plasmid (Addgene #158221) (10) following the manufacturer’s instructions, as described previously (11). Three days after transfection, the cell suspension was centrifuged at 300 × g for 5 min to remove cells from the conditioned medium. For Expi293F-palmGRET EV separation, the conditioned medium was centrifuged at 2,000 × g for 20 min, filtered through a 0.22 μm bottle-top system (Corning), and concentrated 10X by tangential flow filtration (TFF, Vivaflow^®^ 50R TFF cassettes, Sartorius). Samples were further concentrated 4X using ultrafiltration (Centricon Plus 70 Ultracel® PL-100, Merck Millipore; 4,000 × g, 20 min, RT) and eluted (1,500 × g, 2 min). Samples were separated into 11 fractions using size-exclusion chromatography (SEC; qEV10 70nm columns, IZON). EV-enriched fractions (1-4) were pooled and concentrated using Amicon 15 Ultra RC 10 kDa MWCO filters (Merck Millipore). Expi293F-palmGRET EV aliquots in Dulbecco’s phosphate buffered saline (DPBS) were stored at 4 ºC for short-term use and at -80 ºC for long-term storage.

### Western blotting

Expi293F-palmGRET cells were harvested and washed twice in DPBS (2,500 × g, 5 min, RT). The resulting pellet was lysed with RIPA buffer (Cell Signaling Technology #9806) for 1 h on ice, vortexing every 15 minutes. The cell lysate was centrifuged at 14,000 × g for 15 min at 4 ºC to remove debris, and the supernatant was quantified using BCA protein assay (Thermo Scientific #23227). EV samples and SEC fractions (10 μl) were lysed with RIPA buffer for 10 min at room temperature. Samples were mixed with reducing (Thermo Scientific #39000) or non-reducing (Thermo Scientific #39001) sample buffer, boiled at 100 ºC for 5 min, and loaded into Criterion TGX stain-free gels (4-15%; Bio-Rad #5678084) with a Spectra™ Multicolor Broad Range Protein Ladder (Thermo Fisher #26634). Gels ran in Tris/Glycine/SDS buffer (Bio-Rad #1610772) at constant voltage (100V) for 1.5 h, and stain-free gels were imaged on the Gel Doc EZ imager (Bio-Rad). Proteins were transferred to PVDF membranes (iBlot™ 2 Transfer Stacks, Invitrogen #IB24001) on the iBlot™ 2 Gel Transfer Device (Invitrogen) using a stacked program (20 V for 1 min; 23 V for 4 min, and 25 V for 2 min). Membranes were blocked with 5% milk-PBST for at least 1 h at room temperature under agitation and incubated with primary antibodies overnight at 4 ºC with orbital shaking. Membranes were probed for CD63 (1:3000, Ms, BD #556016), CD9 (1:3000, Ms, BioLegend #312102), syntenin (1:1000, Rb, Abcam #133267) and calnexin (1:1000, Rb, Abcam #22595), washed four times with 5% milk-PBST for 5 min under agitation, and exposed to the appropriate secondary antibody (m-IgGκ BP-HRP, 1:5000, Santa Cruz #616102 or Mouse anti-Rb IgG HRP, 1:5000, Santa Cruz #2327) for 1 h, RT under agitation. Membranes were washed four times with PBST, exposed for 60 seconds to SuperSignal™ West Pico ECL substrate (Thermo Scientific #34578), and documented using the iBright 1500 (Invitrogen).

### Flow nanoanalyzer

Expi293F-palmGRET EV preparations were measured for particle concentration, size distribution, and EGFP incidence (%) using a Flow NanoAnalyzer (NanoFCM Co., Ltd). System start-up included laser alignment and calibration with fluorescent 250 nm silica nanoparticles (2.19E+10 particles/ml, NanoFCM #QS2503) for particle concentration and a premixed silica nanosphere cocktail with monodisperse nanoparticle populations of 68 nm, 91 nm, 113 nm, and 155 nm in diameter (NanoFCM #516M-Exo) for size dispersion. Blank was set with DPBS as the vehicle. EV samples were diluted 1:1000 (v/v) in DPBS and boosted for 60 seconds before particle signal acquisition, using constant pressure of 1 kPa and event rates between 1,500 and 10,000 events/min. Signals for side-scattering and EGFP signal, particle concentration, and size dispersion were calculated using NanoFCM Professional Suite V2.0 software.

### Transmission electron microscopy

Expi293F-palmGRET EV stock was diluted 1:20 (v/v) and adsorbed to glow-discharged carbon-coated 400 mesh copper grids by flotation for 2 min. Grids were quickly blotted and rinsed by floatation on three drops of Tris-buffered saline, 1 min each, before being negatively stained in two consecutive drops of 1% uranyl acetate (UAT) with tylose (1% UAT in deionized water (diH2O), double filtered through a 0.22-μm filter). Grids were blotted and quickly aspirated to cover the sample with a thin layer of stain. Grids were imaged on a Hitachi 7600 transmission electron microscope (TEM) operating at 80 kV with an AMT XR80 CCD (8 megapixel), under magnification ranging from 30,000x to 120,000x.

### Coverglass chamber preparation for EV immobilization

Schott D 263^®^ coverglass chambers (μ-Slide 8 Well high Glass Bottom, ibidi #80807) were coated with 300 μl of poly-L-lysine solution (0.1% PLL in H_2_O (w/v), Sigma #P8920) and incubated overnight at 4 ºC. The coating residue was removed through two washes with 500 μl of DPBS (Gibco). Expi293F-palmGRET EVs (4E+08 particles/ml in DPBS, 300 μl per well) were used to coat both PLL-coated and uncoated chambers and were incubated overnight at 4 ºC and protected from light. The next day, wells were washed with DPBS twice (300 μl per well) and filled with either 300 μl of DPBS or 100 μl of VECTASHIELD® mounting medium (Vector Laboratories #H-1000-10). Samples were stored at 4 ºC until analysis. Three independent repeats with technical duplicates were done.

### Coverslip preparation for EV immobilization

Borosilicate glass coverslips (Carolina Cover Glasses, Circle 12 mm, Thickness 0.13-0.17 mm, Carolina Science & Math #NC9537307) and quartz coverslips (Ø10mm × 0.25mm thick, Ted Pella #26010) were rinsed with 70% ethanol and dried before use. Placed atop a 30 μl drop of EV dilution (4E+08 particles/ml) on parafilm, coverslips were incubated overnight at 4 ºC, protected from light. Coverslips were rinsed twice with DPBS and placed onto clear histological slides using VECTASHIELD^®^ mounting medium. Slides were sealed (CoverGrip Coverslip Sealant, Biotium #23005) and stored at 4 ºC until analysis. The experiment was conducted with two slips per group, in three independent repeats, using two distinct EV preparations.

### Direct stochastic optical reconstruction microscopy (dSTORM)

A super-resolution microscopy system (Nanoimager, Oxford NanoImaging), coupled with NimOS software, was calibrated using beads detected using 405/473/532/635 nm laser excitation, mounted atop a 100x oil-objective lens. Good quality channel mapping calibration was obtained through adequate point coverage. Coverslips and coverglass chambers were imaged in seven random sites (79.28 μm × 49.43 μm) with 1000 frames at 30 ms exposure and 1.4 numerical aperture. Laser intensity was optimized for each sample. Post-acquisition spatial analysis was performed using CODI (alto.codi.bio), using the “clustering and counting” app for correcting drift, clustering populations, and counting particles. Widefield images were acquired under 1x zoom, and reconstituted images were snapped acquired under 6-8x zoom.

### B cell isolation

Non-human primate samples were obtained from SPF juvenile male pigtailed macaques *(Macaca nemestrina)* in procedures approved by the Johns Hopkins University Animal Care and Use Committee (ACUC) and conducted following the Weatherall Report, the Guide for the Care and Use of Laboratory Animals, and the USDA Animal Welfare Act. Blood samples were diluted (1:1, v/v) in Hank’s balanced salt solution (HBSS) and applied to SepMate™-15 (IVD) tubes loaded with a density gradient (Percoll®, Cytiva #17-0891-01). Blood was spun at 1,200 × g for 10 min for separation into three phases, and the upper fraction, containing mononuclear cells (MNC), was transferred to fresh tubes and washed with HBSS. Cells were pelleted (300 × g for 8 min) and treated with red blood cell lysis buffer (ACK lysing buffer 0.83% NH_4_Cl, 0.1% KHCO_3_, 0.03% EDTA) for 10 min at 37 ºC. Cells were washed with HBSS, spun at 300 × g for 8 min, and resuspended in cold selection buffer (PBS + 0.5% BSA + 2 mM EDTA) mixed with 20% (v/v) CD20 MicroBeads (Miltenyi Biotec #130-091-104 and #130-042-201) for 15 min at 4 ºC. The suspension was diluted in selection buffer, centrifuged at 300 × g for 8 min, resuspended in buffer, and applied onto MS columns (Miltenyi Biotec #130-042-201) for magnetic cell sorting using MiniMACS™ separators (Miltenyi Biotec). CD20^+^ B cells were collected in conical tubes, pelleted (300 × g for 8 min), and resuspended in RPMI 1640 + 10% FBS + 1% pen/strep for cell counting.

### B cell adhesion to EV coating

Coverglass chambers (μ-Slide 8 Well high Glass Bottom, ibidi #80807) were coated with 300 μl poly-L-lysine (PLL, 0.1%, v/v) for 2 h at 4 ºC and washed with DPBS (300 μl/well). An EV suspension (4E+08 particles/ml in DPBS) was used to coat wells overnight at 4 ºC, protected from light. Wells coated with 1% BSA-PBS (200 μl/well) were used as the negative control. Excess coating was washed off with 300 μl DPBS, and non-specific binding was blocked with 300 μl 1% BSA-PBS in each well, followed by a 1-hour incubation at 37 ºC. B cells (1E+05 cells in 300 μl RPMI 1640 10% FBS) were applied to each well and incubated at 37 ºC, 5% CO_2_, for 1 h. Unbound cells were removed with three washes with DPBS (300 μl per well) before fixation with 4% PFA for 10 min at RT. Cells were washed twice with PBS, permeabilized with 0.1% Triton X-100 for 5 min at RT, and washed again. F-actin was stained with Phalloidin-647 (200 μl/well, 1:1000 dilution in PBS, Abcam #176759) for 1 h in the dark, and excess dye was washed off twice with PBS. Wells were coated with Prolong Mounting Medium with DAPI and sealed with parafilm for analysis using the Nanoimager (ONI). Cell count was assessed via nuclei count using FIJI (ImageJ), imaging a minimum of 6 random sites under 20x magnification using a confocal microscope (Zeiss 880 Airyscan FAST). The experiment was conducted on two independent occasions, using duplicated wells per group in each repeat.

### Statistical analysis

Raw data were checked for outliers using the ROUT method (12), assuming a Q = 0.1. D’Agostino-Pearson omnibus K2 normality test was applied to identify data distribution. Parametric data were analyzed using unpaired t-test or one-way ANOVA with Tukey’s multiple comparison test, and non-parametric data were analyzed using the Mann-Whitney test. Statistical analysis and graphs were executed on GraphPad Prism (v. 10.0.1 (170)).

## Results

### Expi293F-palmGRET EV characterization

Expi293F cells were cultured to 720 ml of culture. Approximately 2.8E+9 cells were transfected with pLenti-palmGRET EGFP-nanoluciferase reporter plasmid. Transfection efficiency was estimated under a fluorescent microscope equipped with a 488nm filter on the first and third days of transfection (Fig. 1A). Cells reached a population of approximately 3.8E+9 cells after three days, at which time EV-containing conditioned culture medium was harvested and EVs were separated. Following the updated MISEV2023 guidelines (13), Expi293F-palmGRET EVs were characterized using multiple techniques. Volumetric Western blotting for proteins CD9, CD63, syntenin, and calnexin indicated the expected depletion of ER-specific calnexin and enrichment of EV-related syntenin and tetraspanins CD63 and CD9 in the early SEC fractions (Fig. 1B). Later SEC fractions were protein-enriched compared with EV fractions (Fig. 1C). EV-enriched fractions (1-4) were pooled for nanoflow analysis, revealing concentration of 4.7E+11 particles/ml, with 78.7% of particles being EGFP^+^ (Fig. 1D). EV size dispersion peaked at 76 nm (Fig. 1E). EVs were visualized with transmission electron microscopy (Fig. 1F) and super-resolution microscopy (Fig. 1G), showing expected size, morphology, and fluorescence.

**Fig 1.**
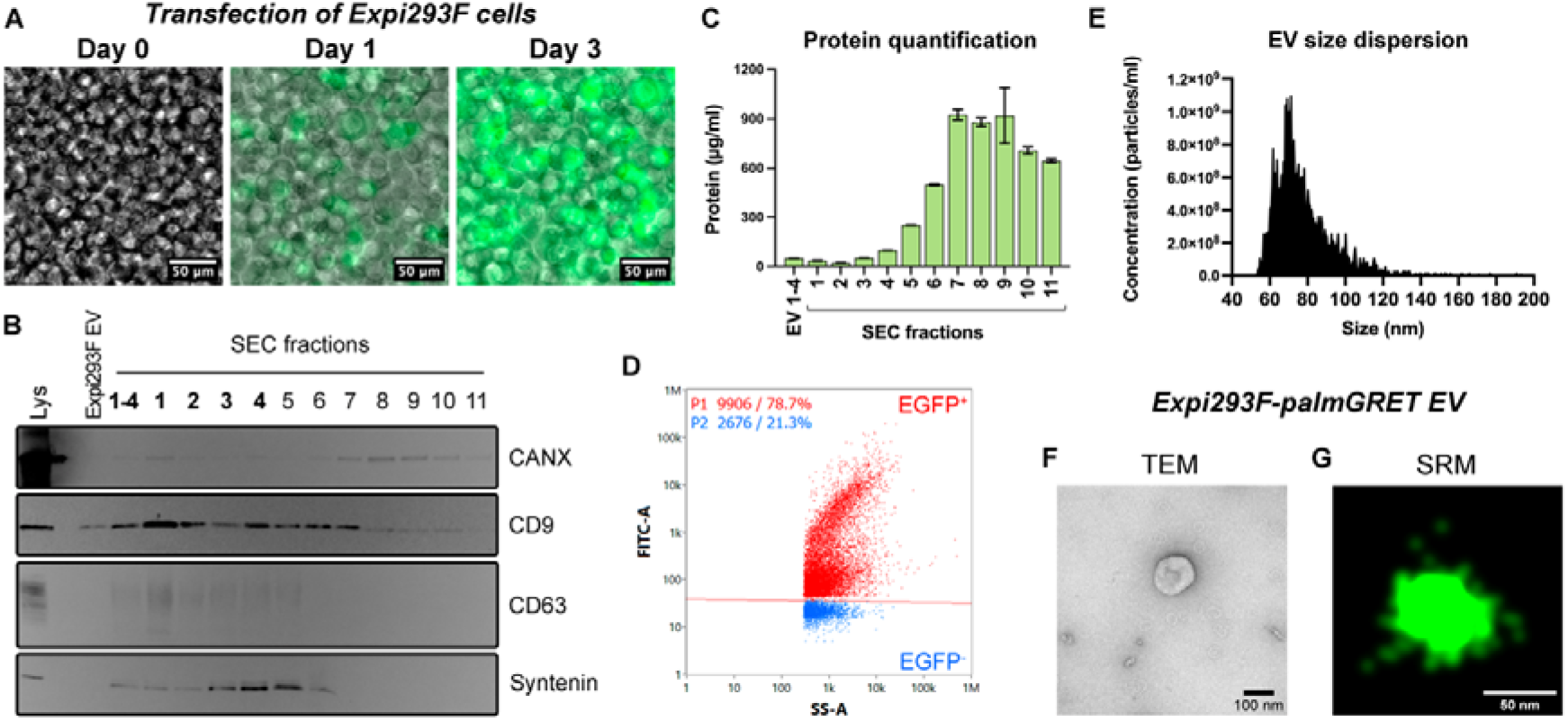
Expi293F-palmGRET EV characterization. (A) Fluorescence imaging of Expi293F cells tracking the three days of transient transfection with pLenti-palmGRET. Scale bar: 50 μm. (B) Western blotting: membranes with Expi293F cell lysate and SEC fractions 1-11 were probed for calnexin (CANX), CD9, CD63, and Syntenin-1. (C) Quantification of protein content in SEC fractions 1-11. (D) Scatter plot of EV nanoflow cytometry, showing 78.7% EGFP-positivity amongst particles from the pooled SEC fractions 1-4. (E) Size distribution plot of Expi293F-palmGRET EVs by nanoflow cytometry. (F) Transmission electron microscopy. Scale bar: 100 nm. (G) Super-resolution microscopy image of a single EV particle emitting EGFP fluorescence. Scale bar: 50 nm.

### EV immobilization is more efficient on glass than on quartz coverslips

Using the protocol depicted in Figure 2A, we found that both quartz and borosilicate glass can immobilize EVs. To count particles, each coverslip was imaged in at least six random sites in dSTORM, and the CODI software clustered signals from the multiple frames to identify individual EVs. Particle count detected 59.62 ± 17.67 on the quartz coverslip and 289.7 ± 172.3 clusters on the borosilicate glass coverslip, indicating a statistically significant difference in EV immobilization (*p* < 0.0001) (Fig. 2B). Inspection of the rendered images indicates the density of particles per frame (Fig. 2C).

**Fig 2.**
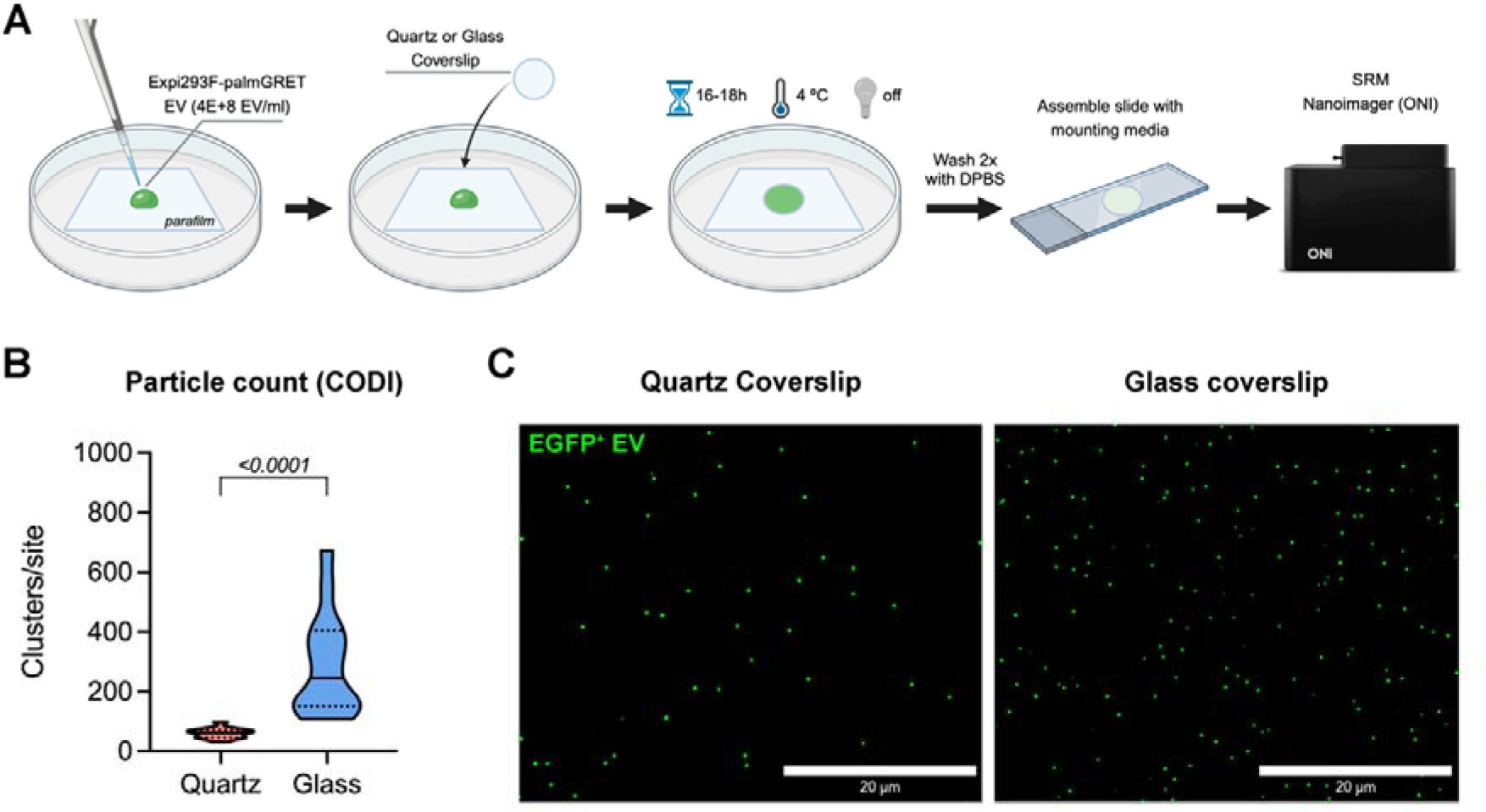
EV immobilization on glass and quartz coverslips. (A) Experimental scheme for the preparation of EV-coated coverslips for SRM imaging. (B) Comparison of particle count between quartz and glass coverslips, obtained using the CODI software (alto.codi.bio). Non-parametric data was compared using the Mann-Whitney unpaired t-test, and the *p*-value is displayed on the graph. (C) Representative images of rendered dSTORM frames, showing individual Expi293F-palmGRET EVs in green, obtained at 60x magnification and 2x zoom. Scale bar: 20 μm.

### EV immobilization is more uniform with PLL pre-coating

Using an 8-well culture chamber assembled atop a Schott D 263 borosilicate coverslip, we cross-examined the requirement of specific mounting medium and PLL to improve EV immobilization on glass. The experimental design for this assay is shown in Figure 3A. Regarding the requirement of mounting medium to stabilize the sample for imaging in SRM, we found a statistically significant difference between PBS *versus* mounting medium (MM) in glass surfaces only, but not in PLL-coated glass (Fig. 3B). This suggests PBS may be a viable medium for immersing samples in culture chambers for dSTORM, if used alongside PLL coating. We also found that EVs attached to borosilicate glass even without PLL coating (no PLL = 157.3 ± 144.9 clusters per site; PLL = 78.83 ± 45.26 clusters per site). However, although there were more EVs in the no PLL condition, high standard deviation (ranging from >50 to 300 particles per site) suggests that PLL facilitates a more uniform immobilization of EVs onto glass. Representative images of the rendered frames indicating the density of particles per site are shown in Fig. 3C.

**Fig 3.**
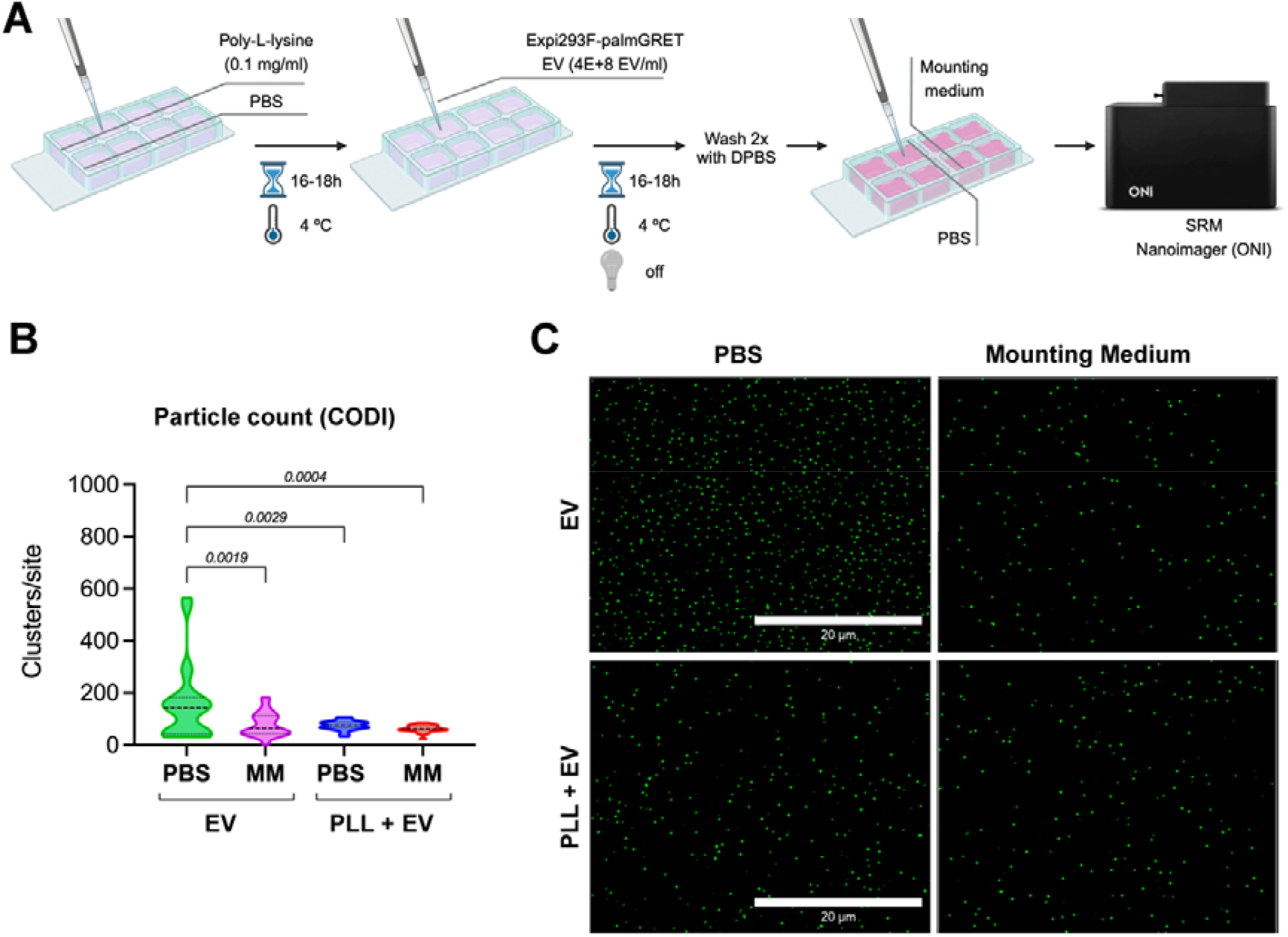
EV immobilization onto borosilicate chamber coverglass. (A) Experimental scheme for the preparation of EV-coated chamber coverglasses for SRM imaging. (B) Comparison of particle count between uncoated wells or pre-coated with poly-L-lysine (0.1 mg/ml) prior to Expi293F-palmGRET EV immobilization and immersed in PBS or mounting medium (MM), obtained using the CODI software (alto.codi.bio). Non-parametric data were compared using the Kruskal-Wallis analysis of variance with Dunn’s multiple comparison test, and the *p*-values are displayed on the graph. (C) Representative images of rendered dSTORM frames, showing individual Expi293F-palmGRET EVs in green, obtained at 60x magnification and 2x zoom. Scale bar: 20 μm.

### B cells interact with PLL/glass-immobilized EVs

We next performed a cell adhesion assay in which B-cells were allowed to interact with an EV-coated coverslip following a previous report (8). For this, we used PLL-coated glass coverslips and non-human primate (NHP) primary CD20^+^ B cells (Fig. 4A). An EV concentration of 4E+8 particles/ml was chosen from a dilution series (Fig. 4B) for subsequent comparison with a negative control (1% BSA-PBS). Although some cells bound to the negative control wells, on average 3.3-fold more cells adhered to EV-coated wells (Fig. 4C); binding of cells was also confirmed by SRM (Fig. 4D).

**Fig 4.**
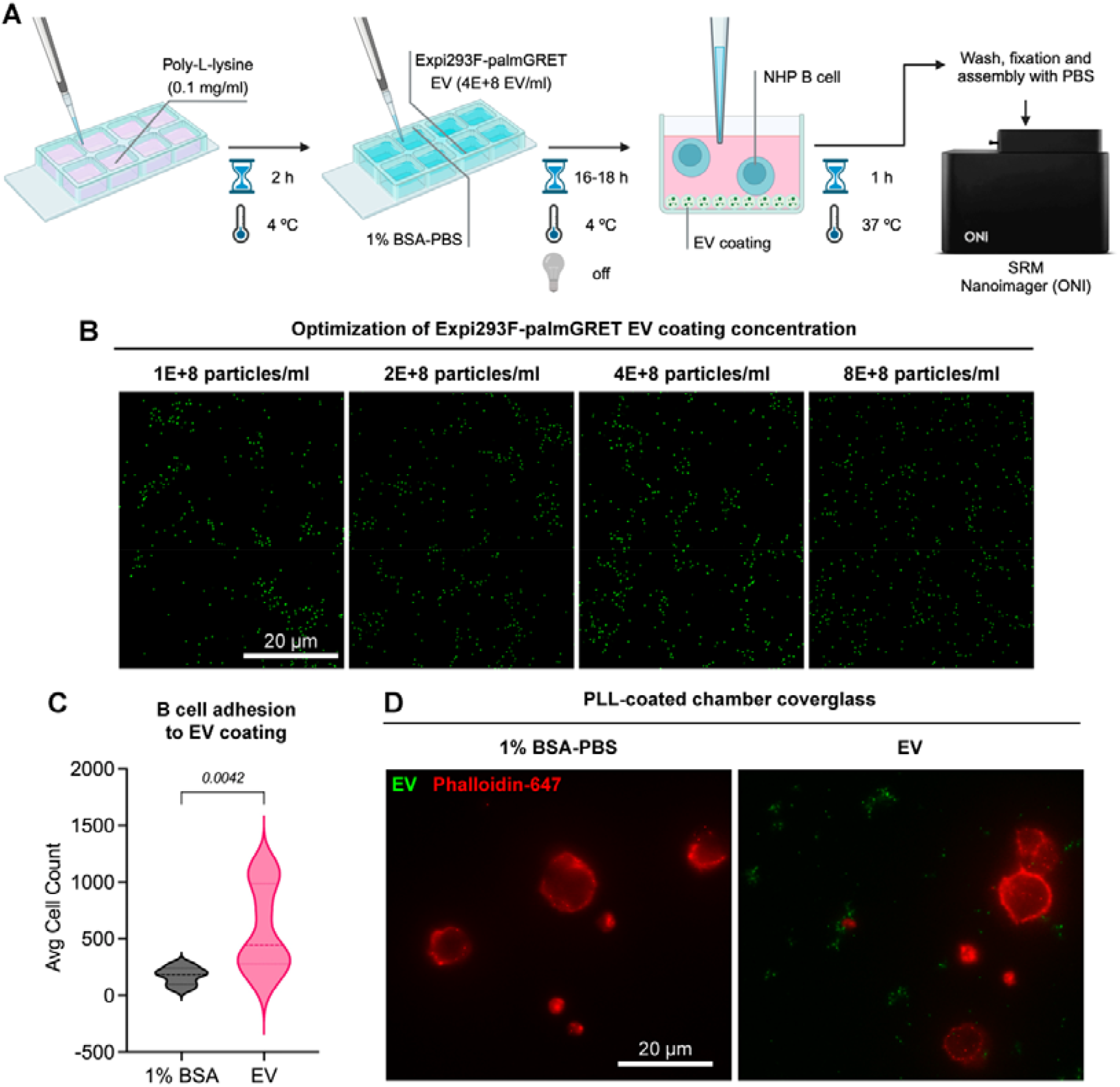
B-cells adhere to EV coating. (A) Experimental scheme. (B) Optimization of EV coating concentration, ranging from 1E+8 to 8E+8 particles/ml. Representative images from SRM rendering indicate Expi293F-palmGRET EVs in green and were obtained at 60x magnification and 2x zoom. Scale bar: 20 μm. (C) Average count of attached cells in 1% BSA-PBS (negative control) and EV coating after 1h of incubation, obtained through nuclei counting. Non-parametric data was compared using the Mann-Whitney unpaired t-test and the *p*-value is displayed on the graph. (D) Representative SRM images from the adhesion assay, showing EVs (green) and NHP B cells (red). Scale bar: 20 μm.

## Discussion

Recently, a paper demonstrated that EV immobilization occurs similarly in quartz glass slides and positively charged quartz slides while describing a novel plugin for particle count using standard fluorescent microscopy images (9). We tested if similar results could be found between standard borosilicate glass coverslips and quartz coverslips, but instead using dSTORM to assess particle disposition. Using EGFP^+^ EVs, a significantly higher mobilization of EVs in glass coverslips rather than quartz was observed, suggesting that the use of commercially available glass coverslips is possible for assays in which EVs need to be adsorbed by an amorphous surface. We also found that the use of PLL coating in borosilicate glass is not required but aids the uniformity of EV distribution. We believe the PLL coating may contribute to EV binding in sites where the glass surface is not as homogeneous, by creating an electrostatic potential to improve EV adhesion (14). On a side note, we also found that mounting medium is not essential for SRM analysis, and PBS can be used as stable media to store microscopy samples under immersion for short periods. To the best of our knowledge, no studies so far have investigated whether PLL is required for EV immobilization on glass surfaces, and few have studied the interaction of EVs with different materials for functional assays.

Sample preparation for specific applications such as total internal reflection fluorescence (TIRF) microscopy and dSTORM often require refined materials that improve their optical transmittance without compromising the roughness value on the surface or the chemical resistance (14). Quartz is considered a superior material for optical transmission, however, its high softening point (1,500 ºC) creates a more delicate material and a costlier product (15). A commercially available option is the standard glass coverslip, made from high-quality borosilicate glass with different sizes and thicknesses (e.g., 0.09 to 0.25 mm), which are suitable for microscopy, especially when treated with PLL in preparation for cell adhesion (16). EVs bind more efficiently to borosilicate glass than quartz, as found using dSTORM, suggesting that this material may be used for the development of new products focused on highly specific microscopy techniques.

Our findings suggest that the immobilization of EVs for highly specific imaging techniques may be achieved by using standard lab materials such as borosilicate glass coverslips. This simpler approach to dSTORM broadens the possibilities for methods and applications that can employ SRM. Also, coupled with the similarity between PBS and mounting medium to store the sample, we described a cheaper option for SRM preparation with equivalent results to costlier options.

## Author’s contribution

BCP: conceptualization, data curation, formal analysis, investigation, methodology, writing – original draft, writing – review and editing; BC: methodology, visualization; SEQ: methodology, visualization; HSSA: funding acquisition, supervision, writing – review and editing; KWW: conceptualization, funding acquisition, supervision, resources, writing – review and editing.

## Acknowledgments

The authors thank members of the Witwer Laboratory, particularly Dr. Olesia Gobololova for consulting; the Retrovirus Laboratory; and the Biochemistry and Molecular Biology Laboratory for helpful suggestions and discussions.

## Data availability

We have submitted all relevant data of our experiments to the EV-TRACK knowledgebase (EV-TRACK ID: EV240144) (17). The data that support the findings of this study are available from the corresponding author upon reasonable request.

